# First detected incursions of avian influenza H5N1 clade 2.3.4.4b into mainland Australia from the Southern Ocean

**DOI:** 10.64898/2026.08.08.743700

**Authors:** Matthew J. Neave, Sam Hair, Patrick Mileto, Jackie E. Mahar, Vittoria Stevens, Kelly Davies, Mark O’Dea, Sadia Iqbal, Jamie W.L. Ong, Andrew Hughes, Jianning Wang, Nathan Fox, Jodie C. Crowder, Donna Gillies, Jeff Butler, Jo Grimsey, Amy McMahon, Michelle Gagliardi, Ebony Grech, Mark Ford, Carlie Soul, Megan Poon, Tristan Reid, Axel Colling, Julie C. McInnes, Tristan L. Burgess, Jarrod C. Hodgson, Thierry Boulinier, David T. Williams, Jasmina M. Luczo, Vidya Bhardwaj, Dwane O’Brien, Debbie Eagles, Guy Baele, Frank Y.K. Wong

## Abstract

High pathogenicity avian influenza H5N1 clade 2.3.4.4b has caused a panzootic of devastating impact to poultry and wildlife globally. The Australian continent and broader Oceania until recently remained the last major region without confirmed detections. Here we report the first H5N1 clade 2.3.4.4b detections from two live seabirds - a brown skua and a southern giant petrel - found on the south coast of Western Australia in June 2026. Virus genome sequencing showed that both viruses were most closely related to H5N1 viruses detected recently on sub-Antarctic islands in the Southern Indian Ocean. In time-calibrated phylogeographic analyses, both viruses sampled in Western Australia clustered with viruses from Heard Island, a sub-Antarctic external territory of Australia. Ancestral location reconstruction also identified Heard Island as the most probable source location, although unsampled intermediate locations cannot be excluded. The two Western Australian detections were estimated to be independent incursions from Heard Island, rather than local transmission on mainland Australia. There was no evidence of reassortment with endemic avian influenza viruses in Australia, and both virus sequences retained key avian-like genetic markers and lacked known substitutions for reduced antiviral susceptibility. These detections revealed a Southern Ocean pathway of recurrent H5N1 incursions into Australia, highlighting the risk of potential establishment on the mainland and the need for heightened surveillance and rapid, nationally-coordinated, virus genomic characterisation.

## Main

High pathogenicity avian influenza (HPAI) H5N1 clade 2.3.4.4b has caused unprecedented mortality in wild birds, poultry and marine mammals in impacted regions globally. After reaching South America in late 2022, the virus entered the Antarctic and Southern Ocean regions, with detections in South Georgia and the Falkland Islands in 2023, Antarctica, the Crozet and Kerguelen Islands in 2024, and Heard Island in 2025 (Banyard et al., 2024; Clessin et al., 2025, 2026; McInnes et al., 2026; Ogrzewalska et al., 2026). Although the precise pathways of virus movement through the Southern Ocean are not fully understood, evidence to date supports a process of repeated circumpolar introduction events (Banyard et al., 2024; Clessin et al., 2026). The remote nature of this region has meant that potential intermediate infection source locations or hosts are largely unsampled. Potential dispersal by wide-ranging pelagic seabirds including skuas, giant petrels, gulls and albatrosses has been suggested (Banyard et al., 2024; Clessin et al., 2026; Wille et al., 2026). The apparent eastward progression of the virus through sub-Antarctic islands established a possible southern Indian Ocean circumpolar incursion route for H5N1 to reach the Australian continent, likely following the Southern Ocean Flyway (Morten et al., 2025). Heard Island was recently confirmed as the first affected Australian external territory (McInnes et al., 2026). Despite these encroaching sub-Antarctic incursions, mainland Australia and the broader Oceania region remained free of HPAI H5N1 clade 2.3.4.4b until June 2026. Here we report the first detections of HPAI H5N1 virus on mainland Australia and Oceania, and present phylogeographic analyses of the virus genome sequences to infer their likely source and route of incursion.

The first H5N1 detection was in a brown skua (*Stercorarius antarcticus lonnbergi*) found in a lethargic and ataxic condition at Cape Le Grand National Park near Esperance, on the remote south coast of Western Australia on June 14^th^ 2026, followed by a second virus detection in a sick (lethargy and hindlimb ataxia) juvenile southern giant petrel (*Macronectes giganteus*) found separately in the same area on June 18^th^ 2026 (Fig. 1). Species level identification of the giant petrel was confirmed by mitochondrial cytochrome *b* barcoding, with the sequence matching the common southern giant petrel genotype “sgp1” (Techow et al., 2010). Real-time reverse transcription PCR testing and influenza A virus sequencing confirmed the detection of HPAI H5N1 clade 2.3.4.4b and infectious virus was isolated from each case.

**Fig. 1.**
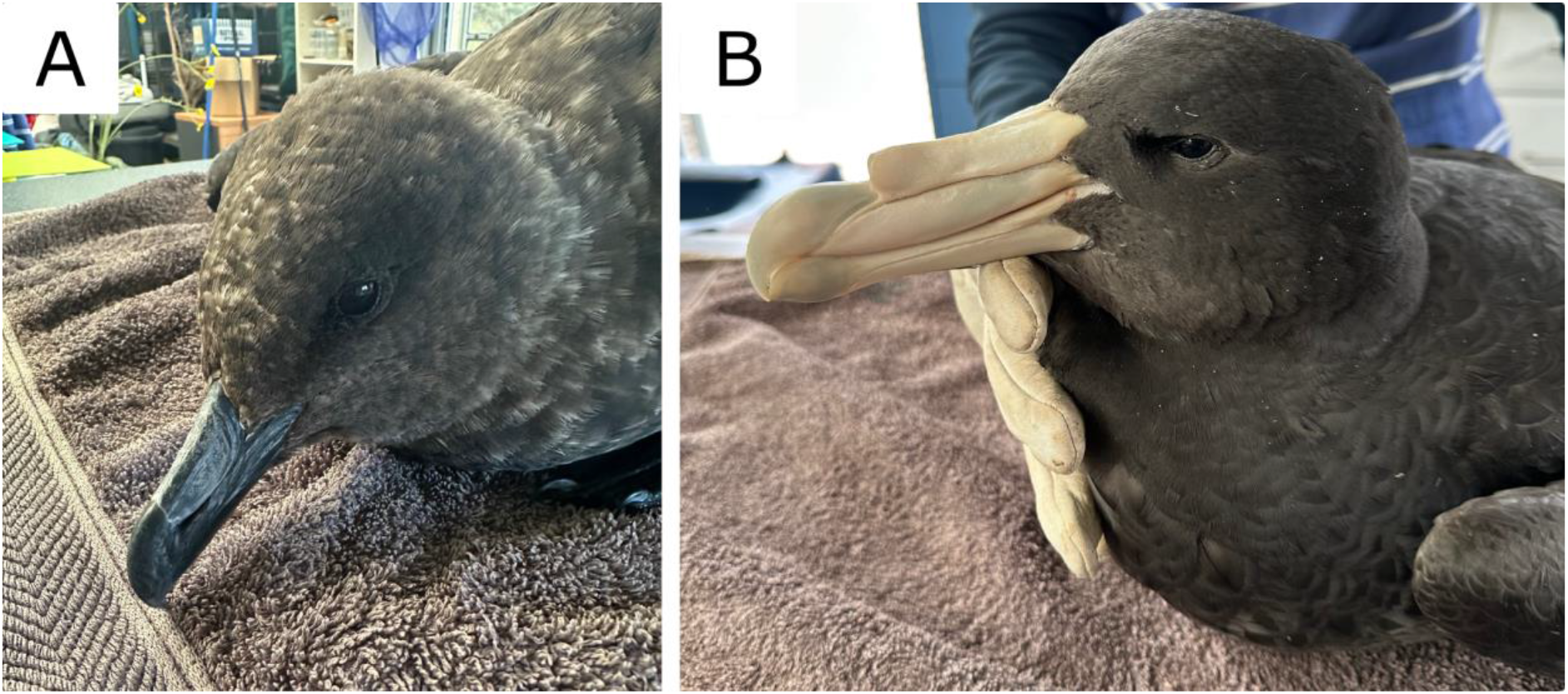
**A** Brown skua (*Stercorarius antarcticus lonnbergi*) and **B** Southern giant petrel (*Macronectes giganteus*) found near Esperance in Western Australia infected with HPAI H5N1 2.3.4.4b avian influenza in June 2026. Photo: L-A Shibish, Esperance Wildlife Sanctuary.

To estimate the origin of the Western Australian H5N1 detections, we analysed their complete virus genomes together with representative H5N1 clade 2.3.4.4b sequences from the Antarctic Peninsula, sub-Antarctic islands and global reference datasets. Time-resolved phylogeographic analyses of the concatenated virus genomes showed that both Western Australian virus samples clustered with H5N1 viruses detected on Heard Island in late 2025 and early 2026 (Fig. 2). A broader Southern Ocean (Heard/Crozet/Kerguelen) radiation was dated to approximately March 2024 (95% HPD: Jan - May 2024), consistent with previous estimates (Clessin et al., 2025, 2026), with Crozet Island virus sequences appearing basal to the Kerguelen and Heard Island infections (McInnes et al., 2026). The inferred most recent common ancestor (MRCA) of the Western Australian brown skua and southern giant petrel infections was estimated to be on Heard Island in approximately August 2025 (95% HPD: Jun - Sep 2025). However, the two samples from Western Australia clustered within different Heard Island sublineages, rather than forming a local Australian cluster (Fig. 2). The brown skua virus sample shared an MRCA around September 2025 (95% HPD: Aug - Oct 2025) with viruses from southern elephant seals (*Mirounga leonina*) sampled on the western side of Heard Island, whereas the southern giant petrel virus had an MRCA of around October 2025 (95% HPD: Sep - Oct 2025) shared with a southern elephant seal virus from the southeast of Heard Island (McInnes et al., 2026). These MRCA nodes had strong support (posterior probability > 0.97), and discrete phylogeographic reconstruction strongly supported Heard Island as the most probable sampled ancestral location of the viruses detected on mainland Australia (Bayes factor = 2,368), which is consistent with progressive easterly circumpolar spread within the sub-Antarctic outbreak region (Clessin et al., 2025, 2026; McInnes et al., 2026). However, the time gap between the inferred MRCA node dates and the June 2026 detections on mainland Australia suggests unsampled ongoing virus circulation on Heard Island or in other undetected Southern Ocean locations before introduction to the Western Australian coast. Our results indicate that the brown skua and southern giant petrel detections represent independent incursions into Australia via discrete transmission chains, rather than lateral transmission from a single source introduction.

**Fig. 2.**
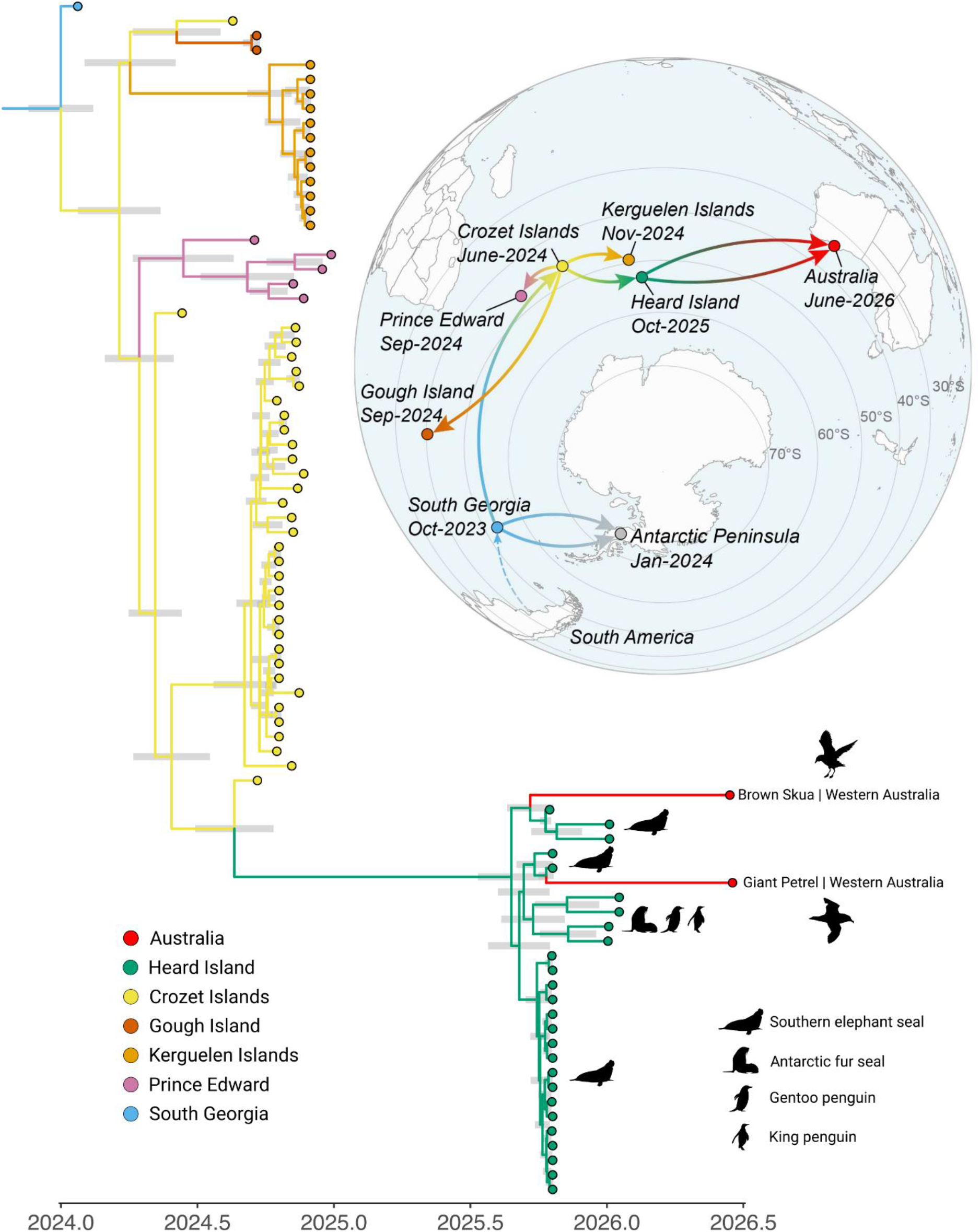
Location-annotated consensus phylogeny of HPAI H5N1 2.3.4.4b spatiotemporal reconstruction across sub-Antarctic islands and Australia shows circumpolar spread into Australia. Only the clade containing Australian, Antarctic and sub-Antarctic sequences (i.e., Clade I; Clessin et al., 2026) is shown for relevance. The likely source of the two Australian introductions was estimated to be Heard Island, with >99% ancestral location support, based on currently available data. Node bars on the internal nodes indicate 95% highest posterior density (HPD) intervals of estimated node ages. Dates on the map are the first reported detection of H5N1 clade 2.3.4.4b at each location. Dots on the map, tip points and external branches are coloured by collection location, while internal branches are coloured by inferred location. Refer to Supplementary Fig. 2 for full tip labels and posterior support values.

Individual gene segment analyses supported the interpretation of independent incursions from the sub-Antarctic islands (Supplementary Fig. 1). Across all eight virus genome segments, the Western Australian viruses were most similar to Heard Island or sub-Antarctic viruses, with nucleotide identities of 99.65-100% to the closest available sequences; while pairwise identities between the two Western Australian viruses were 99.40-99.93% (Supplementary Table 1). Notably, several virus gene segments from the Western Australian samples were identical to segments from Heard Island southern elephant seal samples, including the brown skua HA segment, which was identical to that from sample HIMI-543, and the southern giant petrel NA segment, which was identical to that from sample HIMI-564 (McInnes et al., 2026). Both virus samples from Western Australia were assigned to H5 clade 2.3.4.4b genotype B3.2 (GenoFLU) and E.2 (ggFLU), which is the genotype associated with all available sub-Antarctic samples. There was no evidence of virus segment reassortment with local Australian lineage avian influenza viruses. The virus gene phylogenies and genotype assignments indicate incursions of complete sub-Antarctic B3.2/E.2 genome constellations, rather than reassortants generated after arrival in Australia.

The avian host species and locations of the detections are consistent with the phylodynamic inference of sub-Antarctic H5N1 clade 2.3.4.4b incursions into mainland Australia. Both brown skuas and southern giant petrels are wide-roaming scavengers and predators that are known to interact with vertebrate colonies on land and at sea, where they opportunistically feed on carcasses (Clessin et al. 2026). Both species are Southern Ocean migratory seabirds known to frequent southern Australian waters (Bonnet-Lebrun et al., 2026; van den Hoff 2011), and both are recognised as potentially important hosts in the epidemiological dynamics of H5N1 in the Southern Ocean. Giant petrels have been among the earliest affected species in Southern Ocean H5N1 outbreaks (e.g., Bennison et al., 2024), and their scavenging behaviour and long-distance movements among sub-Antarctic islands and adjacent waters make them plausible mediators of viral spread (Clessin et al., 2026, 2025; Iervolino et al., 2026). Skuas have been prominent in H5N1 Antarctic outbreaks, including in the first reported H5N1 mortality event in Antarctica in 2024, and have been proposed as a sentinel species for HPAI due to their high mortality rates and wide distribution (Bennett-Laso et al., 2024; Iervolino et al., 2026; Wille et al., 2026). Notably, brown skuas and southern giant petrels were observed feeding on H5N1 infected seal carcasses on Heard Island in October 2025 and January 2026 (McInnes et al. 2026). These detections provided observational evidence that these Southern Ocean seabirds can potentially distribute H5N1 to other locations, such as Australia.

Sick, weak or dead pelagic birds are known to wash up on beaches for a number of reasons, sometimes due to complex biophysical processes in adjacent oceans (Parrish et al., 2007). Juvenile giant petrels are commonly seen along the Australian coastline during their first 6-12 months, including birds from Heard Island and Crozet (van den Hoff 2011; Voisin 1990). The timing of the detections also coincided with a major Southern Ocean frontal system affecting the sub-Antarctic region and southern Western Australia (Bureau of Meteorology, 2026).

Virus genome analysis of the Western Australian H5N1 samples showed that they did not possess key mutations known to reduce susceptibility to antivirals including oseltamivir, baloxavir or adamantanes, retaining residues I117, E119, Q136, I223, S247, H275 and N295 in the neuraminidase (NA), polymerase acidic (PA) I38, and matrix (M2) S31 (Suttie et al., 2019). Both virus samples retained avian-like genetic markers, including polymerase basic (PB2) residues E627 and D701. Several substitutions previously described in Heard Island and other sub-Antarctic island H5N1 viruses were also present, including PA N383D, nucleoprotein (NP) I41V, and haemagglutinin (HA1) S133A/S154N/T156A and HA2 K64E (Clessin et al., 2026; McInnes et al., 2026), further supporting their relationship to the southern Indian Ocean outbreak lineage.

Infectious virus was isolated from both Western Australian samples in specific pathogen-free (SPF) embryonated chicken eggs and antigenically characterised by haemagglutination inhibition (HI) assay using post-infection prime chicken antisera raised to representative H5 clade 2.3.4 and 2.3.2 viruses (Supplementary Table 2). The isolates showed highest antigenic reactivity to antisera raised to contemporary clade 2.3.4.4b viruses, including A/bovine/Ohio/B24OSU-439/2024(H5N1) and A/duck/Pampanga/2022(H5N1), consistent with their phylogenetic placement.

## Conclusion

Our findings confirm that mainland Australia has been exposed to separate incursions of H5N1 clade 2.3.4.4b from Southern Ocean migratory seabirds but provide no evidence for sustained transmission on mainland Australia or reassortment with other lineages of low pathogenicity avian influenza viruses that circulate in Australia, based on currently available data. The detection of two infected seabirds from a remote coastline likely represents only a small subset of recent incursion events, given the surveillance challenges and logistical difficulty of detecting sick or dead pelagic birds on long stretches of sometimes remote coastline and the common occurrence of Southern Ocean birds washing ashore, especially in their first year. The occurrence of repeated independent incursions suggests that potential establishment in Australian wildlife is increasingly likely if exposure continues. Ongoing Southern Ocean and Australian wildlife surveillance, virus genome sequencing and integration of pathogen genomic, ecological and meteorological data is essential to assessing the impacts of future incursions and potential ongoing intracontinental transmission. The finding of two infected wide-ranging scavenging seabird species epidemiologically linked to the infections on Heard Island suggests that this threat has been moving further east and could soon impact further sub-Antarctic territories of the Southern Pacific Ocean.

## Methods

### Sample Collection

On June 14^th^, a brown skua (*Stercorarius antarcticus lonnbergi*) was found sick on a beach near Esperance, Western Australia, and was taken to the Esperance Wildlife Hospital. Choanal and cloacal swabs were collected and placed in viral transport medium (VTM) and submitted for testing at the Western Australian Department of Primary Industries and Regional Development (DPIRD) and forwarded to the CSIRO Australian Centre for Disease Preparedness (ACDP) for confirmatory testing. On June 18^th^, a southern giant petrel (*Macronectes giganteus*) was also found sick near Esperance, and tracheal and cloacal swabs were again submitted to DPIRD and then forwarded to ACDP.

### Real-Time Reverse-Transcription PCR

Nucleic acid was extracted from the swabs using the MagMAX-96 viral RNA isolation kit (Thermo Fisher Scientific) according to the manufacturer’s instructions. The swabs were tested for influenza A virus using real-time RT-PCR assays targeting the influenza A matrix (M) gene and the H5 haemagglutinin (HA) gene. The influenza A matrix gene assay was a modified method based on Spackman et al. (2002). The assay used one forward primer, IVA D161 M 5’-AGATGAGYCTTCTAACCGAGGTCG, four reverse primers: IVA D162 M1 5’-GCAAAAACATCYTCAAGTCTCTG, IVA D162 M2 5’-TGCAAACACATCYTCAAGTCTCTG, IVA D162 M3 5’-TGCAAAGACATCYTCAAGTCTCTG, and IVA D162 M4 5’-TGCAAATACATCYTCAAGTCTCTG, and probe IVA M 5’-FAM-TCAGGCCCC/ZEN/CTCAAAGCCGA-IBFQ. The H5 assay was a duplex real-time RT-PCR targeting both the N-terminal and C-terminal regions of the HA gene. The N-terminal assay used forward primer IVA D204F 5’-ATGGCTCCTCGGRAACCC, reverse primers IVA D205R 5’-TTYTCCACTATGTAAGACCATTCCG, with probe IVA D215P 5’-FAM-ATGTGTGACGAATTCMT-MGBNFQ. The C-terminal assay used primers IVA D148 H5 5’-AAACAGAGAGGAAATAAGTGGAGTAAAATT and IVA D149 H5 5’-AYCCRTTSGAGCACATCCA, with probe IVA H5a 5’-FAM-TCAACAGTGGCGAGTTCCCTAGCA-IBFQ. All real-time RT-PCR assays were performed using the AgPath One-Step RT-PCR Kit (Thermo Fisher Scientific) in a final reaction volume of 25 μL. Thermal cycling conditions consisted of reverse transcription at 45°C for 10 min, initial denaturation at 95°C for 10 min, followed by 45 cycles of 95°C for 15 s and 60°C for 45 s.

### Host Barcoding

To confirm and clarify the morphological identification of the birds, host barcoding markers were sequenced from the nucleic acid extract used above. A fragment of mitochondrial cytochrome c oxidase subunit I (COI) was amplified using the conserved avian primers AWCF1 and AWCR6 (Patel et al., 2010) using the HotStarTaq Master Mix Kit (Qiagen). PCR cycling conditions for COI amplification were 94°C for 15 mins, followed by 35 cycles of 94°C for 30 s, 49°C for 45 s, 72°C for 60 s, with a final extension of 72°C for 5 min. For additional species resolution of the giant petrel, a fragment of mitochondrial cytochrome *b* was amplified using the primers GPcytbF and GPcytbR (Techow et al., 2010) using the same PCR kit. PCR cycling conditions were 94°C for 2 mins, followed by 30 cycles of 94°C for 45 s, 55°C for 45 s, 72°C for 60 s, with a final extension of 72°C for 5 min. Capillary sequencing reactions for both amplicons were performed using Big Dye X-terminator (Applied Biosystems) and sequenced using a 3500 xL Genetic Analyzer (Applied Biosystems).

### Viral Isolation and haemagglutination inhibition assay

The VTM swab samples were inoculated into 9-11 day old embryonated specific pathogen-free chicken eggs (3 eggs per sample), and the eggs incubated at 37°C. Eggs were candled twice daily to examine for the presence of a live embryo. As embryos either approached the humane end point and were chilled, or were found dead, the allantoic fluid was harvested and subjected to a haemagglutination assay to confirm the presence of live virus, and virus sequencing used to confirm isolation of H5N1 virus. The virus isolation in embryonated chicken eggs was conducted under approved application ACDP23009, approved by the CSIRO – ACDP Animal Ethics Committee. The antigenic relationship of the virus isolates was investigated by haemagglutination inhibition (HI) assay using a panel of chicken antisera raised against bovine and poultry HPAI H5 viruses belonging to clade 2.3.4 and 2.3.2. HI assays were performed with chicken red blood cells using standard techniques (WOAH, 2024).

### Whole-genome sequencing

Viral RNA was extracted from the swabs using the MagMAX-96 viral RNA isolation kit (Thermo Fisher Scientific) according to the manufacturer’s instructions. All influenza A genome segments were amplified using the SuperScript III one-step RT-PCR system with high-fidelity Platinum Taq DNA polymerase (Thermo Fisher Scientific) and universal influenza A virus primers: MBTuni-12 5’-ACGCGTGATCAGCAAAAGCAGG and MBTuni-13 5’-ACGCGTGATCAGTAGAAACAAGG as previously described (Kampmann et al., 2011; Zhou et al., 2009). The amplified products were processed using the Oxford Nanopore Rapid Barcoding Kit (SQK-RBK114.24) and sequenced on R10 MinION flow cells. Sequence reads were trimmed for quality and mapped to reference influenza A sequences using Minimap2 v.2.24 (Li 2018) within the Geneious Prime software v.2026.0.1 (Biomatters, Auckland, NZ). In addition, the cleaned reads were assembled using IRMA v.1.0.2 (Shepard et al., 2016). The final consensus genomes were manually curated and annotated using Geneious Prime. The genomes were screened for molecular markers of mammalian adaptation, altered virulence and antiviral resistance using FluMut v.0.6.4 (Giussani et al., 2025).

### Phylogenetic analysis

All publicly available H5N1 clade 2.3.4.4b HA genes with complete sampling dates as of June 1st, 2026, were downloaded from GISAID (gisaid.org) and complemented with the HA genes of the two Western Australian genomes generated as part of this study. The HA genes were aligned using MAFFT v.7.490 (Katoh and Standley, 2013) as part of the nextclade pipeline (Aksamentov et al., 2021), subsequently trimmed and checked for outliers using root-to-tip regression in TempEst (Rambaut et al., 2016). A maximum-likelihood phylogenetic tree of the resulting 14,163 HA sequences was estimated using IQ-TREE v.3.1.3 (Nguyen et al., 2015), placing the two Western Australian sequences together with sequences from the Crozet and Kerguelen Islands, Gough and Marion Island, and the Antarctic Peninsula (data not shown).

Given this determination of a plausible sub-Antarctic origin for the H5N1 introductions into mainland Australia, the backbone data sets described by Clessin et al. (2026) and McInnes et al. (2026) were subsequently used to combine publicly available genomes from GISAID and BV-BRC (bv-brc.org) with 26 Heard Island genomes (McInnes et al., 2026), 21 genomes from the Crozet and Kerguelen Islands (Clessin et al., 2025, 2026), and the two Western Australian genomes generated here. The final dataset comprised 1,324 genomes. The individual segments were first aligned using MAFFT v.7.490 (Katoh and Standley, 2013) and maximum-likelihood phylogenetic trees were estimated using IQ-TREE v.3.1.3 (Nguyen et al., 2015). Segment alignments were then concatenated and an ML tree generated to increase phylogenetic resolution. As the interpretation was the same for the concatenated tree and the HA tree (and segments with appropriate resolution), the concatenated alignment was subsequently used for the time-structured analyses.

Time-calibrated phylogenetic analysis was performed on the concatenated genome alignment using Bayesian inference through Markov chain Monte Carlo as implemented in BEAST X v.10.5.0 (Baele et al., 2025b). The analyses assumed a GTR+Γ substitution model, an uncorrelated relaxed molecular clock with an underlying lognormal distribution (Drummond et al., 2006) and a non-parametric skygrid coalescent prior (Gill et al., 2013). Hamiltonian Monte Carlo was used to improve sampling efficiency of the skygrid (Baele et al., 2020) and molecular clock parameters (Ji et al., 2020). Default BEAST X priors were used. Four independent analysis replicates were run for 500 million iterations with GPU acceleration through the BEAGLE v4 library (Gangavarapu et al., 2026), then combined after removal of 10% burn-in. Convergence and mixing were assessed in Tracer v.1.7.2 (Rambaut et al., 2018) with all ESS values > 200. Majority-rule highest independent posterior subtree reconstruction (HIPSTR) was used to generate consensus trees in TreeAnnotator X (v.10.5.0) (Baele et al., 2025a).

### Phylogeographic analysis

We performed discrete phylogeographic inference using both asymmetric forward-in-time (FIT; Lemey et al., 2009) and backward-in-time (BIT; De Maio et al., 2015; Shao et al., 2026) continuous-time Markov chain models on the Clade I subtree (Clessin et al., 2026) using BEAST X (Baele et al., 2025b) with BEAGLE v4 support (Gangavarapu et al., 2026). Bayesian stochastic search variable selection was employed for FIT and BIT models, and default BEAST X priors were used. The analyses were run for 250 million iterations after which convergence and mixing were assessed in Tracer v.1.7.2 (Rambaut et al., 2018) with all ESS values > 200. Majority-rule HIPSTR was used to generate consensus trees in TreeAnnotator X (v.10.5.0) (Baele et al., 2025a).

## Acknowledgments

We gratefully acknowledge all data contributors, i.e., the authors and their originating laboratories responsible for obtaining the specimens, and their submitting laboratories for generating the genetic sequence and metadata and sharing via the GISAID initiative, on which this research is based. T.B. acknowledges support from French Polar Institute (IPEV ECOPATH-1151), ANR (WILDFLU ANR-25-CE35-0691) and CNRS (EE, SEE-Life).

G.B. acknowledges support from the Research Foundation – Flanders (“Fonds voor Wetenschappelijk Onderzoek – Vlaanderen,” G098321N), from the European Union Horizon 2023 RIA project LEAPS (grant agreement no. 101094685), and from the DURABLE EU4Health project 02/2023-01/2027 which is co-funded by the European Union (call EU4H-2021-PJ4) under Grant Agreement No. 101102733.

## Data availability

Whole-genome sequence data from both viruses has been deposited in GISAID (gisaid.org) under accessions: brown skua (EPI_ISL_20479340) and southern giant petrel (EPI_ISL_20497638).

## References

Aksamentov, I., Roemer, C., Hodcroft, E.B., Neher, R.A. 2021. Nextclade: clade assignment, mutation calling and quality control for viral genomes. Journal of Open Source Software, 6(67), 3773, 10.21105/joss.03773

Baele, G., Gill, M. S., Lemey, P., Suchard, M. A. 2020. Hamiltonian Monte Carlo sampling to estimate past population dynamics using the skygrid coalescent model in a Bayesian phylogenetics framework. Wellcome Open Res. 2020, 5: 53.

Baele, G., Carvalho, L. M., Brusselmans, M., Dudas, G., Ji, X., McCrone, J. T., Lemey, P., Suchard, M. A., Rambaut, A. 2025a. HIPSTR: highest independent posterior subtree reconstruction in TreeAnnotator X. Bioinformatics 41(10): btaf488

Baele, G., Ji, X., Hassler, G.W., McCrone, J.T., Shao, Y., Zhang, Z., Holbrook, A.J., Lemey, P., Drummond, A.J., Rambaut, A., Suchard, M.A., 2025b. BEAST X for Bayesian phylogenetic, phylogeographic and phylodynamic inference. Nat. Methods 22, 1653–1656. 10.1038/s41592-025-02751-x

Banyard, A.C., Bennison, A., Byrne, A.M.P., Reid, S.M., Lynton-Jenkins, J.G., Mollett, B., De Silva, D., Peers-Dent, J., Finlayson, K., Hall, R., Blockley, F., Blyth, M., Falchieri, M., Fowler, Z., Fitzcharles, E.M., Brown, I.H., James, J., 2024. Detection and spread of high pathogenicity avian influenza virus H5N1 in the Antarctic Region. Nat. Commun. 15, 7433. 10.1038/s41467-024-51490-8

Bennett-Laso, B., Berazay, B., Muñoz, G., Ariyama, N., Enciso, N., Braun, C., Krüger, L., Barták, M., González-Aravena, M., Neira, V., 2024. Confirmation of highly pathogenic avian influenza H5N1 in skuas, Antarctica 2024. Front. Vet. Sci. 11, 1423404. 10.3389/fvets.2024.1423404

Bennison, A., Adlard, S., Banyard, A.C., Blockley, F., Blyth, M., Browne, E., Day, G., Dunn, M.J., Falchieri, M., Fitzcharles, E., Forcada, J., Forster Davidson, J., Fox, A., Hall, R., Holmes, E., Hughes, K., James, J., Lynton-Jenkins, J., Marshall, S., McKenzie, D., Morley, S.A., Reid, S.M., Stubbs, I., Ratcliffe, N., Phillips, R.A., 2024. A case study of highly pathogenic avian influenza (HPAI) H5N1 at Bird Island, South Georgia: the first documented outbreak in the subantarctic region. Bird Study 71, 380–391. 10.1080/00063657.2024.2396563

Bonnet-Lebrun, A.-S., Delord, K., Cherel, Y., Ribout, C., Guillou, G., Bustamante, P., Barbraud, C., 2026. Post-breeding season behaviour of a threatened population of subtropical brown skuas. Mar. Biol. 173, 49. 10.1007/s00227-025-04770-w

Bureau of Meteorology, 2026. Analysis Chart Archive [WWW Document]. URL https://www.bom.gov.au/australia/charts/archive/ (accessed 6.27.26).

Clessin, A., Briand, F.-X., Tornos, J., Lejeune, M., De Pasquale, C., Fischer, R., Souchaud, F., Hirchaud, E., Hong, S.L., Bralet, T., Guinet, C., McMahon, C.R., Grasland, B., Baele, G., Boulinier, T., 2025. Circumpolar spread of avian influenza H5N1 to southern Indian Ocean islands. Nat. Commun. 16, 8463. 10.1038/s41467-025-64297-y

Clessin, A., Brusselmans, M., Hong, S.L., Tornos, J., Lejeune, M., Shao, Y., Briand, F.-X., Abolnik, C., Kaza, B., Suchard, M.A., Aguado, B., Alcamí, A., Barbraud, C., Beer, M., Bennison, A., Bonadonna, F., Bonnet, T., Bost, C.-A., Boucheron, S., Bralet, T., Catry, P., Cleeland, J., Connan, M., Coombes, H.A., Delord, K., De Pasquale, C., Dewar, M.L., Dong, X., Emerit, J., Fischer, R., Fountain-Jones, N., Galimberti, F., González-Solís, J., Guinet, C., Gunn, C., Günther, A., Iervolino, M., James, J., Ji, X., Jonsen, I., Jones, C.W., Kuepfer, A., Kuiken, T., Lebohec, C., Lisovski, S., Zubiri, L.L., Lynton-Jenkins, J.G., Martinez-García, P., McCulley, M., McMahon, C.R., Mollett, B.C., Moraga-Quintanilla, A.I., Nichol, R., Noiret, A., Ogrzewalska, M., Owen, K., Pardo-Roa, C., Peroteau, S., Phillips, R.A., Poulin, E., Rambaut, A., Reid, S.M., Riehle, E., Risi, M., Rumianowski, O., Ryan, P.G., Sanvito, S., Stanworth, A., Steinfurth, A., Stevenson, J., Stier, A., Uhart, M.M., Vanstreels, R.E.T., Vázquez-Calvo, A., Vianna, J.A., Wells, K., White, J., Whitelaw, P., Wille, M., Younger, J., Roberts, L.C., Grasland, B., Banyard, A.C., Nelson, M.I., Gamble, A., Boulinier, T., Baele, G., 2026. Dispersal, adaptation and persistence of H5N1 in the sub-Antarctic and Antarctica. 10.64898/2026.03.20.713283

De Maio, N., Wu, C.-H., O’Reilly, K.M., Wilson, D., 2015. New Routes to Phylogeography: A Bayesian Structured Coalescent Approximation. PLoS Genet 11, e1005421. 10.1371/journal.pgen.1005421

Drummond, A.J., Ho, S.Y.W., Phillips, M.J., Rambaut, A., 2006. Relaxed phylogenetic and dating with confidence. PLOS Biol. 4(5): e88

Gangavarapu, K., Ji, X., Shao, Y., Lemey, P., Rambaut, A., Baele, G., Suchard, M.A., 2026. BEAGLE 4.1: A high-performance library for computation on phylogenetic trees across diverse parallel architectures. arXiv:2606.27607v1 10.48550/arXiv.2606.27607

Gill, M.S., Lemey, P., Faria, N.R., Rambaut, A., Shapiro, B., Suchard, M.A., 2013. Improving Bayesian Population Dynamics Inference: A Coalescent-Based Model for Multiple Loci. Molecular Biology and Evolution 30, 713–724. 10.1093/molbev/mss265

Giussani, E., Sartori, A., Salomoni, A., Cavicchio, L., De Battisti, C., Pastori, A., Varotto, M., Zecchin, B., Hughes, J., Monne, I., Fusaro, A., 2025. FluMut: a tool for mutation surveillance in highly pathogenic H5N1 genomes. Virus Evol. 11, veaf011. 10.1093/ve/veaf011

Iervolino, M., Günther, A., Begeman, L., Aguado, B., Bestebroer, T.M., Bellido-Martin, B., Coerper, A., Fornillo, M.V., Fusaro, B., Ibañez, A.E., Leijten, L., Lisovski, S., Mañez, M.B., Reade, A., Van Run, P., Soto, F., Wallis, B., Dewar, M., Alcamí, A., Beer, M., Vanstreels, R.E.T., Kuiken, T., 2026. The expanding H5N1 avian influenza panzootic causes high mortality of skuas in Antarctica. Sci. Rep. 16, 4604. 10.1038/s41598-025-34736-3

Ji, X., Zhang, Z., Holbrook, A., Nishimura, A., Baele, G., Rambaut, A., Lemey, P., Suchard, M. A. 2020. Gradients do grow on trees: a linear-time O(N)-dimensional gradient for statistical phylogenetics. Mol. Biol. Evol. 37(10): 3047–3060.

Kampmann, M.-L., Fordyce, S.L., Ávila-Arcos, M.C., Rasmussen, M., Willerslev, E., Nielsen, L.P., Gilbert, M.T.P., 2011. A simple method for the parallel deep sequencing of full influenza A genomes. J. Virol. Methods 178, 243–248. 10.1016/j.jviromet.2011.09.001

Katoh, K., Standley, D.M., 2013. MAFFT Multiple Sequence Alignment Software Version 7: Improvements in Performance and Usability. Mol. Biol. Evol. 30, 772–780. 10.1093/molbev/mst010

Lemey, P., Rambaut, A., Drummond, A.J., Suchard, M.A., 2009. Bayesian Phylogeography Finds Its Roots. PLoS Comput Biol 5, e1000520. 10.1371/journal.pcbi.1000520

Li, H., 2018. Minimap2: pairwise alignment for nucleotide sequences. Bioinformatics 34, 3094–3100. 10.1093/bioinformatics/bty191

McInnes, J.C., Burgess, T., Mergard, G., Wells, M.R., McMahon, C.R., Neave, M.J., Polanowski, A., Terauds, A., Tornos, J., Lejeune, M., Briand, F.-X., Baele, G., Boulinier, T., Achurch, H., Alderman, R., Lashko, A., Wienecke, B., Wynen, L.P., Viola, B., Virtue, P., Hodgson, J.C., 2026. Mass mortality of southern elephant seals during multi-species outbreak of HPAI H5N1 on sub-Antarctic Heard Island. 10.64898/2026.06.16.732752

Morten, J.M., Carneiro, A.P.B., Beal, M., Bonnet‐Lebrun, A., Dias, M.P., Rouyer, M., Harrison, A., González‐Solís, J., Jones, V.R., Garcia Alonso, V.A., Antolos, M., Arata, J.A., Barbraud, C., Bell, E.A., Bell, M., Bose, S., Broni, S., De L Brooke, M., Butchart, S.H.M., Carlile, N., Catry, P., Catry, T., Charteris, M., Cherel, Y., Clark, B.L., Clay, T.A., Cole, N.C., Conners, M.G., Debski, I., Delord, K., Egevang, C., Elliot, G., Esefeld, J., Facer, C., Fayet, A.L., Fijn, R.C., Fischer, J.H., Franklin, K.A., Gilg, O., Gill, J.A., Granadeiro, J.P., Guilford, T., Handley, J.M., Hanssen, S.A., Hawkes, L.A., Hedd, A., Jaeger, A., Jones, C.G., Jones, C.W., Kopp, M., Krietsch, J., Landers, T.J., Lang, J., Le Corre, M., Mallory, M.L., Masello, J.F., Maxwell, S.M., Medrano, F., Militão, T., Millar, C.D., Moe, B., Montevecchi, W.A., Navarro‐Herrero, L., Neves, V.C., Nicholls, D.G., Nicoll, M.A.C., Norris, K., O’Dwyer, T.W., Parker, G.C., Peter, H., Phillips, R.A., Quillfeldt, P., Ramos, J.A., Ramos, R., Rayner, M.J., Rexer‐Huber, K., Ronconi, R.A., Ruhomaun, K., Ryan, P.G., Sagar, P.M., Saldanha, S., Schmidt, N.M., Schultz, H., Shaffer, S.A., Stenhouse, I.J., Takahashi, A., Tatayah, V., Taylor, G.A., Thompson, D.R., Thompson, T., Van Bemmelen, R., Vicente‐Sastre, D., Vigfúsdottir, F., Walker, K.J., Watts, J., Weimerskirch, H., Yamamoto, T., Davies, T.E., 2025. Global Marine Flyways Identified for Long‐Distance Migrating Seabirds From Tracking Data. Global Ecol Biogeogr 34, e70004. 10.1111/geb.70004

Nguyen, L.-T., Schmidt, H.A., Von Haeseler, A., Minh, B.Q., 2015. IQ-TREE: A Fast and Effective Stochastic Algorithm for Estimating Maximum-Likelihood Phylogenies. Mol. Biol. Evol. 32, 268–274. 10.1093/molbev/msu300

Ogrzewalska, M., Vanstreels, R.E.T., Pereira, E.C., Campinas, E., Correa Junior, L., Melo, J.O., Macedo, L., Appolinario, L.R., Arantes, I., Brandao, M.L., Pribul, B.R., Rocha, C.M., Campos, F.S., Souza, U.J.B.D., Galarza, B.P., Melgarejo, A.S., Trilles, L., Moreira, L.M., Degrave, W., Magalhães, M., Moreira, D., Vilela, R.D.V., Motta, F.C., Siqueira, M.M., Resende, P.C., 2026. Genomic analysis of high pathogenicity avian influenza viruses from Antarctica reveals multiple introductions from South America. Nat. Commun. 17, 4927. 10.1038/s41467-026-71544-3

Parrish, J., Bond, N., Nevins, H., Mantua, N., Loeffel, R., Peterson, W., Harvey, J., 2007. Beached birds and physical forcing in the California Current System. Mar. Ecol. Prog. Ser. 352, 275–288. 10.3354/meps07077

Patel, S., Waugh, J., Millar, C.D., Lambert, D.M., 2010. Conserved primers for DNA barcoding historical and modern samples from New Zealand and Antarctic birds. Mol. Ecol. Resour. 10, 431–438. 10.1111/j.1755-0998.2009.02793.x

Rambaut, A., Lam, T.T., Carvalho, L.M., Pybus, O. 2016. Exploring the temporal structure of heterochronous sequences using TempEst (formerly Path-O-Gen). Virus Evol. 2(1): vew007

Rambaut, A., Drummond, A.J., Xie, D., Baele, G., Suchard, M.A., 2018. Posterior Summarization in Bayesian Phylogenetics Using Tracer 1.7. Syst. Biol. 67, 901–904. 10.1093/sysbio/syy032

Shao, Y., Suchard, M.A., Rambaut, A., Ji, X., Lemey, P., Vasylyeva, T.I., Baele, G. 2026. Parallel algorithms for phylogenetic inference under a structured coalescent approximation. Proc. Natl. Acad. Sci. USA 123 (18) e2602412123

Shepard, S.S., Meno, S., Bahl, J., Wilson, M.M., Barnes, J., Neuhaus, E., 2016. Viral deep sequencing needs an adaptive approach: IRMA, the iterative refinement meta-assembler. BMC Genomics 17, 708. 10.1186/s12864-016-3030-6

Spackman, E., Senne, D.A., Myers, T.J., Bulaga, L.L., Garber, L.P., Perdue, M.L., Lohman, K., Daum, L.T., Suarez, D.L., 2002. Development of a Real-Time Reverse Transcriptase PCR Assay for Type A Influenza Virus and the Avian H5 and H7 Hemagglutinin Subtypes. J Clin Microbiol 40, 3256–3260. 10.1128/JCM.40.9.3256-3260.2002

Suttie, A., Deng, Y.M., Greenhill, A.R., Dussart, P., Horwood, P.F., Karlsson, E.A., 2019. Inventory of molecular markers affecting biological characteristics of avian influenza A viruses. Virus Genes 55(6):739–768. 10.1007/s11262-019-01700-z

Techow, N.M.S.M., O’Ryan, C., Phillips, R.A., Gales, R., Marin, M., Patterson-Fraser, D., Quintana, F., Ritz, M.S., Thompson, D.R., Wanless, R.M., Weimerskirch, H., Ryan, P.G., 2010. Speciation and phylogeography of giant petrels Macronectes. Mol. Phylogenet. Evol. 54, 472–487. 10.1016/j.ympev.2009.09.005

van den Hoff, J., 2011. Recoveries of juvenile Giant Petrels in regions of ocean productivity: potential implications for population change. Ecosphere 2, art75. 10.1890/ES11-00083.1

Voisin, J-F., 1990. Movements of giant petrels Macronectes spp. banded as chicks at Iles Crozet and Kerguelen. Marine ornithology 18, 27–36.

Wille, M., Abbott, W., Day, D., Deng, Y.-M., Dong, X., Gibson, T., Hope-Inglis, R., McCulley, M., Olsson, I., Varsani, A., Visentin, T., Walters, M., Dewar, M., 2026. Skuas as sentinels of high pathogenicity avian influenza H5N1 on the Antarctic Peninsula in the 2024/2025 austral summer. 10.64898/2026.02.15.706047

World Organisation for Animal Health (WOAH). 2024. Chapter 3.3.4. Avian influenza (including infection with high pathogenicity avian influenza viruses). In: Manual of Diagnostic Tests and Vaccines for Terrestrial Animals, 13th edn. WOAH, Paris.

Zhou, B., Donnelly, M.E., Scholes, D.T., St. George, K., Hatta, M., Kawaoka, Y., Wentworth, D.E., 2009. Single-Reaction Genomic Amplification Accelerates Sequencing and Vaccine Production for Classical and Swine Origin Human Influenza A Viruses. J. Virol. 83, 10309–10313. 10.1128/JVI.01109-09

