## Supplementary Figures and Tables for "First detected incursions of avian influenza H5N1 clade 2.3.4.4b into mainland Australia from the Southern Ocean"

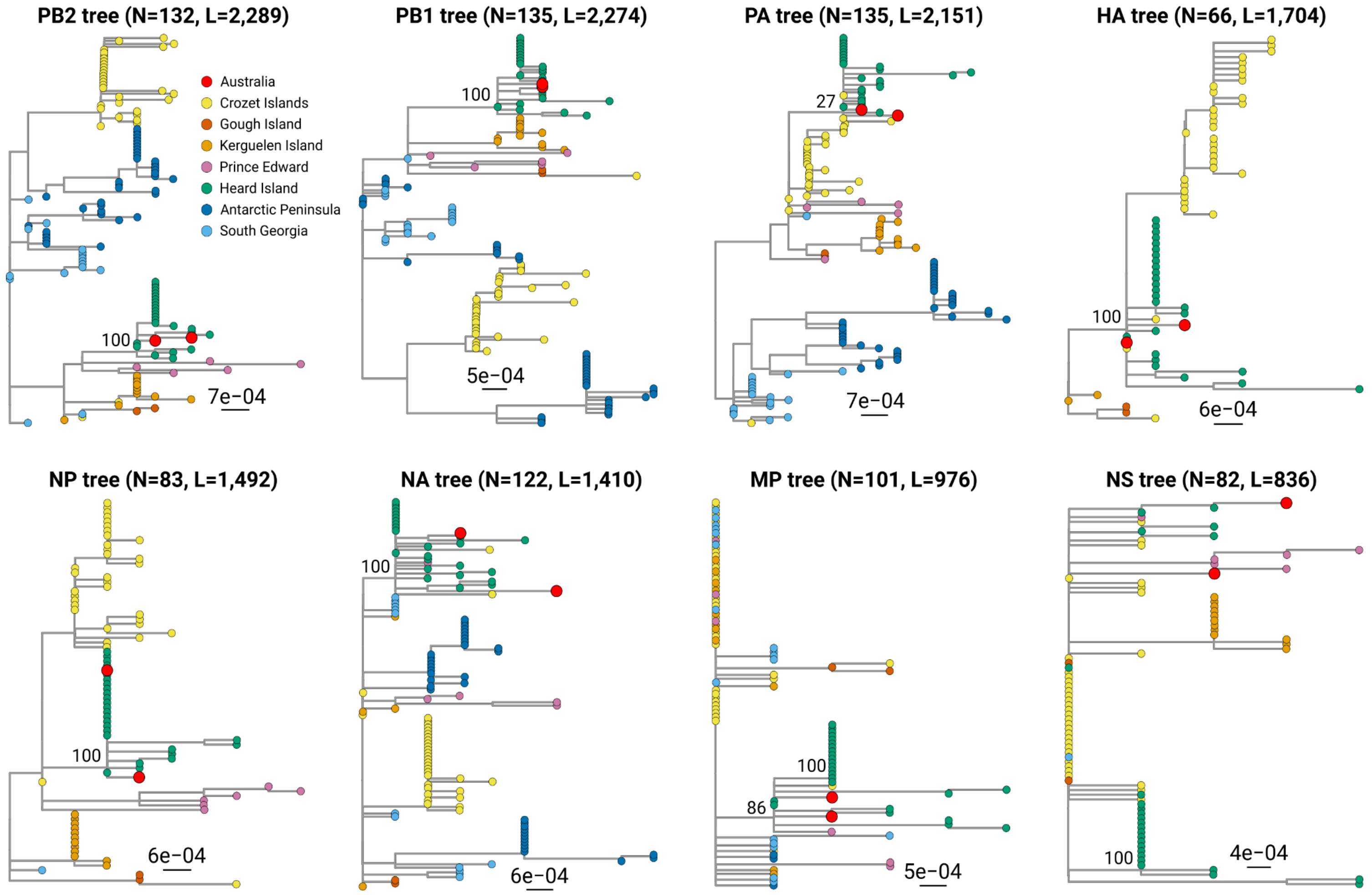


**Supplementary Fig. 1 | Maximum-likelihood phylogenies of each genome segment of the Western Australian and sub-Antarctic HPAI H5N1 clade 2.3.4.4b viruses.** The trees were inferred for each genome segment using the full reference dataset of approximately 1,300 sequences, then visualised by extracting the branch containing the Australian, Heard Island and Crozet Island sequences. The scale bar indicates the average number of substitutions per site. Tip-points are coloured by collection location and support from 1,000 ultrafast bootstraps are given for key nodes. The title of each plot gives the number of included sequences (N) and the nucleotide alignment length (L).

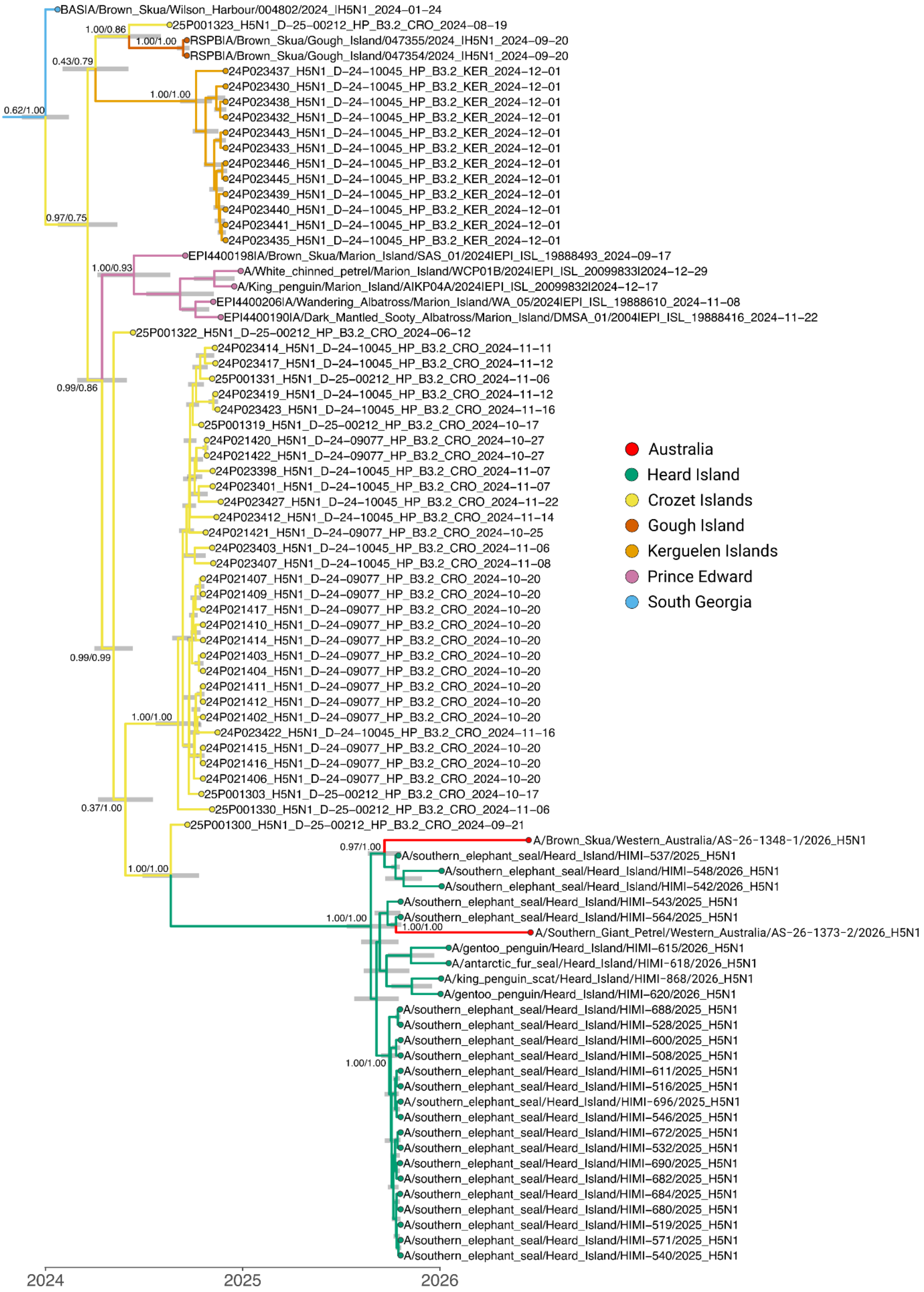


**Supplementary Fig.2 | Location-annotated consensus phylogeny of HPAI H5N1 2.3.4.4b spatiotemporal reconstruction.** Only the branch containing Australian, Antarctic and sub-Antarctic sequences (i.e., Clade I; Clessin et al., 2026) is shown for relevance. Node bars on the internal nodes indicate 95% highest posterior density (HPD) intervals of estimated node ages, and node labels indicate “posterior branch support / ancestral state support”. Tip points and external branches are coloured by collection location, while internal branches are coloured by inferred location.

**Supplementary Table 1 |** Segment-wise percent nucleotide similarity values for each segment of the two Western Australian HPAI H5N1 viruses and the closest matching reference sequence, and pairwise similarity between the Western Australian samples.

| Segment | Brown skua closest match | % ID | Southern giant petrel closest match | % ID | Brown skua vs southern giant petrel %ID |
|---|---|---|---|---|---|
| PB2 | Southern Elephant Seal \| Heard \| HIMI-543 \| 2025-10-21* | 99.87 | Southern Elephant Seal \| Heard \| HIMI-543 \| 2025-10-21* | 99.96 | 99.82 |
| PB1 | Southern Elephant Seal \| Heard \| HIMI-537 \| 2025-10-17 | 99.91 | Southern Elephant Seal \| Heard \| HIMI-564 \| 2025-10-21 | 100.00 | 99.82 |
| PA | Southern Elephant Seal \| Heard \| HIMI-508 \| 2025-10-21* | 99.95 | Southern Elephant Seal \| 24P021411 \| Crozet \| 2024-10-20* | 100.00 | 99.81 |
| HA | Southern Elephant Seal \| Heard \| HIMI-543 \| 2025-10-21 | 100.00 | Southern Elephant Seal \| Heard \| HIMI-543 \| 2025-10-21 | 99.88 | 99.88 |
| NP | Southern Elephant Seal \| Heard \| HIMI-508 \| 2025-10-21* | 100.00 | Southern Elephant Seal \| Heard \| HIMI-508 \| 2025-10-21* | 99.93 | 99.93 |
| NA | Southern Elephant Seal \| Heard \| HIMI-543 \| 2025-10-21* | 99.65 | Southern Elephant Seal \| Heard \| HIMI-564 \| 2025-10-21 | 100.00 | 99.50 |
| MP | Southern Elephant Seal \| Heard \| HIMI-543 \| 2025-10-21* | 99.90 | Southern Elephant Seal \| Heard \| HIMI-543 \| 2025-10-21* | 99.90 | 99.80 |
| NS | Gentoo Penguin \| Heard \| HIMI-615 \| 2026-01-18* | 99.76 | Southern Elephant Seal \| Heard \| HIMI-564 \| 2025-10-21 | 99.88 | 99.40 |

*Multiple best matches

**Supplementary Table 2 |** Antigenic analysis of H5 avian influenza viruses isolated from the Western Australian samples using HI assay and post-infection prime chicken antisera raised to representative H5 high pathogenicity avian influenza reference strains.

| Test Sample No. | A/bovine/Ohio/B24OSU-439/2024 (H5N1) Clade 2.3.4.4b | A/duck/Pampanga/22-0524-1/2022 (H5N1) Clade 2.3.4.4b | A/avian/Nepal/456/2021 (H5N8) Clade 2.3.4.4b | A/avian/Philippines/20-2489-P1/2020 (H5N6) Clade 2.3.4.4e | A/duck/Dili/747/2022 (H5N1) Clade 2.3.2.1g |
|---|---|---|---|---|---|
| A/Brown Skua/Western Australia/AS-26-1348-1/2026 (H5N1) Isolate 1 | 512 | 512 | 64 | 64 | 32 |
| A/Brown Skua/Western Australia/AS-26-1348-2/2026 (H5N1) Isolate 2 | 256 | 256 | 32 | 32 | 16 |
| A/Giant Southern Petrel/Western Australia/AS-26-1373-1/2026 (H5N1) Isolate 1 | 256 | 128 | 16 | 8 | 16 |
| A/Giant Southern Petrel/Western Australia/AS-26-1373-2/2026 (H5N1) Isolate 2 | 256 | 256 | 32 | 16 | 16 |
| Homologous Reference Antigen* | 512 | 1024 | 128 | 512 | 512 |

*HI titres to homologous antigen are shown on the bottom row; shading indicates highest HI titre and closest 'best-fit' match for each isolate.
